# Scarless SARS-CoV-2 Genome Engineering and Variant Analysis

**DOI:** 10.64898/2026.08.21.746147

**Authors:** Agnieszka Dabrowska, Ashley Cuell, Rahul Basu, Jyoti Vishwakarma, Renee Delgado, Emilia Barreto-Duran, Xingyu Liu, Li He, Yan Xiang, Chengjin Ye, Luis Martinez-Sobrido, Reuben S. Harris

## Abstract

In addition to causing cold and flu-like symptoms, Severe Acute Respiratory Syndrome Coronavirus 2 (SARS-CoV-2) can also cause chronic longer-term diseases. Antiviral drugs, especially used combinatorially, have the potential to reduce the severity of individual infections and prevent the development of chronic disease. One of the safest and most versatile reverse genetics systems for SARS-CoV-2 studies is a bacterial artificial chromosome (BAC)-based system harboring the WA1 strain full-length genome and attenuating deletions in the accessory open reading frame 3a and 7b proteins (*ORF*3a and *ORF*7b, respectively). Here, a scarless genome engineering technique called *En Passant* mutagenesis was used to change one amino acid in the viral main protease (M^pro^ P132) into the residue present in contemporary Omicron strains (H132), in order to more accurately study protease inhibitors and resistance mechanisms. This recombinant, attenuated viral system yields antiviral EC_50_ values for the active component of approved drugs including nirmatrelvir (Paxlovid) and ensitrelvir (Xocova) and, importantly, also enables a parallel assessment of drug efflux. For instance, the antiviral potency of nirmatrelvir improves 50-fold by inhibiting the P-Glycoprotein (P-Gp) transporter with ritonavir or tariquidar, whereas the potency of ensitrelvir is unaffected. This system also enables the safe isolation and characterization of viral variants with reduced sensitivity to drugs, as evidenced by M^pro^ M49L compromising the efficacy of ensitrelvir. Together, these systems combine to provide safe, reliable, and quantitative approaches for M^pro^ variant analysis and drug testing without the biosafety concerns of conducting these experiments using wildtype isolates.

**IMPORTANCE:** Safe genetic systems for studying coronavirus biology and developing next generation antivirals are important. One of the most versatile systems leverages a bacterial artificial chromosome to efficiently propagate and engineer a full-length SARS-CoV-2 genome. This system is also safe because it has crippling deletion mutations that limit virus replication to a small number of cell lines. Here, we use a genome engineering technology to change a single amino acid in the viruses’ main protease enzyme to match that of circulating Omicron isolates. The resulting attenuated virus was also used to demonstrate antiviral efficacy of approved drugs and uncover mutants with reduced drug sensitivity. The emergent mutants match those in a subset of circulating strains further demonstrating broad relevance.

## INTRODUCTION

In the wake of the COVID-19 pandemic, Severe Acute Respiratory Syndrome Coronavirus 2 (SARS-CoV-2/SARS2) continues to circulate in the human population. Due to a continuously changing viral spike (S) protein, prior exposure to ancestral viruses and/or vaccine antigens may not prevent future infections, particularly with antigenically distinct variants. Moreover, SARS2 is only one of a much broader family of coronaviruses that includes animal viruses with zoonotic potential (1). The net result is a near-infinite number of genetic combinations that are likely to result in future regional outbreaks like Severe Acute Respiratory Syndrome Coronavirus (SARS-CoV, SARS1) and Middle East Respiratory Syndrome Coronavirus (MERS-CoV), and potentially in a global pandemic like SARS2.

Viral enzymes such as proteases and replicases are targets for antiviral drug development because they are essential for virus replication and pathogenesis, constrained evolutionarily due to multiple functional roles, and under low selective pressure from adaptive immune responses due to their intracellular nature (*i.e*., rarely exposed to neutralizing antibodies and therefore less likely to be selected evolutionarily). For example, the main protease (M^pro^) of SARS2 cleaves the viral polyprotein (pp)1a and pp1ab into functional non-structural proteins (Nsps) essential for viral replication, including M^pro^ itself (*aka*., Nsp5) (2). Accordingly, M^pro^ has been a top target for SARS2 antiviral drug development, leading to drugs including Paxlovid [a combination of the covalent M^pro^ inhibitor nirmatrelvir (NMV) and the P-Glycoprotein (P-Gp) efflux pump inhibitor ritonavir (RTV)] and Xocova [the non-covalent M^pro^ inhibitor ensitrelvir (ESV)] (3, 4). Additional candidate drugs are in varying stages of development and next-generation coronavirus M^pro^ inhibitors are essential to minimize the problem of drug resistance and to be ready for the next local outbreak or global pandemic by exerting broader-spectrum activity (5, 6).

Multiple reverse genetic systems have been developed to study the biology of coronaviruses and test candidate antiviral compounds (7–16). These systems include the ligation of multiple segments *in vitro*, the use of vaccinia virus as a vector, and the leveraging of bacterial artificial chromosomes (BACs). BAC systems have several advantages, including rapid growth and amplification in *E. coli*, safety due to no virus production in bacteria, and relative ease of genetic engineering due to high rates of homologous recombination. To further ensure safety, *ORF3a* and *ORF7b* were deleted from a WA1 SARS2 genome (WT SARS2) in the context of a BAC system (pBeloBAC11-SARS2-*ΔORF3a-ΔORF7b*) (17–19). ORF3a is an accessory protein that suppresses type I interferon signaling by blocking STAT1 phosphorylation and has also been associated with ion channel activity, inflammasome activation, and apoptosis (20–23). ORF7b is a small membrane-associated accessory protein implicated in modulation of host innate immune responses, including suppression of IFN-β induction through the RIG-I/MDA5–MAVS pathway, and has also been linked to apoptotic cell death, although its precise functions remain less well defined than those of ORF3a (24–27). However, these two ORFs are dispensable for virus replication in several cell lines, including Vero-E6-ACE2-TMPRSS2 (Vero-E6-AT), but required for replication and pathogenesis *in vivo* in murine and golden Syrian hamster models (18, 21).

In this study, a scarless genome engineering technique called *En Passant* mutagenesis was used to change the proline (P) 132 codon in the ancestral WA1 *nsp5* gene into a histidine (H) codon in pBeloBAC11-SARS2-*ΔORF3a-ΔORF7b*, in order to better model M^pro^ functionality and druggability of currently circulating Omicron variants. The resulting recombinant attenuated (RA)SARS2-*ΔORF3a-ΔORF7b*-M^pro^-H132 (RA-SARS2) virus exhibits sensitivity to the M^pro^ drugs ESV and NMV. In addition, the potency of NMV, but not ESV, can be boosted 50-fold by simultaneous treatment with the P-Gp inhibitors RTV or tariquidar (TQR), which recapitulate a known liability of NMV. Furthermore, continuous culture experiments reveal multiple independent M^pro^ M49L variants with reduced sensitivity to ESV, which also recapitulate observations in laboratory and clinical settings. Taken together, these studies demonstrate the utility of the RA-SARS2 system in studying Omicron M^pro^ functionality and druggability, overcoming the biosafety concerns of doing these experiments using WT forms of the virus.

## MATERIALS AND METHODS

### Recombinant BAC engineering

The *E. coli* strain of GS1783 containing a recombinant attenuated (RA)SARS2 (WA1 strain) BAC (pBeloBAC11-RA-SARS-CoV-2 ΔORF3a ΔORF7b, RA-SARS2-M^pro^-P132 BAC) was described previously (17, 18). The RA-SARS2-M^pro^-P132 BAC was engineered here using *En Passant* mutagenesis (28, 29) to convert M^pro^-P132 into M^pro^-H132, which reflects contemporary Omicron isoloates. First, the Kan^R^ cassette was amplified from pEPkan-S (gift from Nikolaus Osterrieder; Addgene plasmid 41017) using Phusion™ High-Fidelity DNA Polymerase (Thermo Fisher Scientific, F530L) and primers with homology to the *nsp5* region of SARS2 (forward: 5’-CAATGGTTCACCATCTGGTGTTTACCAATGTGCTATGAGGCACAATTTCACTATTAAGGGTTCAGGATGACGACGATAAGTAGGG; reverse: 5’-CACATGAACCATTAAGGAATGAACCCTTAATAGTGAAATTGTGCCTCATAGCACATTGGTAAACAACCAATTAACCAATTCTGATTAG). Next, the resulting 1128 bp PCR product containing *nsp5*-flanking homology, a single I-SceI cleavage site, and the Kan^R^ cassette was gel-purified using a GeneJET Gel Extraction Kit, (Thermo Fisher Scientific, K0692) and quantified using a NanoDrop One^C^ UV-Vis Spectrophotometer (Thermo Fisher Scientific, 13400518). GS1783 with the RA-SARS2-M^pro^-P132 BAC was then grown at 32°C to mid-log phase (OD_600_ 0.5 to 0.7) in Luria broth (LB) supplemented with 34 µg/mL chloramphenicol, heat shocked at 42°C for 15 min in a shaking water bath to activate Red-mediated recombination, chilled on ice for 1 hr, and washed 3 times in ice-cold PBS. The cells were then electrotransformed with 500 ng of the Kan^R^ PCR product (electroporation was performed using a Bio-Rad Gene Pulser Xcell system with settings of 1.5 kV, 25 µF, and 200 Ω). After, each electrotransformed reaction was allowed 1.5 hrs of recovery in LB medium at 32°C and then plated on LB plates containing 34 µg/mL chloramphenicol (Goldbio, C-105-25) and 30 µg/mL kanamycin (Goldbio, K-120-25). A PCR strategy was then used to identify single colonies with the Kan^R^ cassette integrated homologously into the *nsp5* region of the RA-SARS2-M^pro^-P132 BAC (Kan-cassette forward primer 5’-TGGTTCACCATCTGGTGTTTAC and reverse primer 5’-AGCCGTTTCTGTAATGAAGGAG, *nsp5*-specific forward primer 5’-TTCTTGGTACAGGCTGGTAATG and reverse primer 5’-TGAGCAGAAAGAGGTCCTAGT).

Next, the Kan^R^ cassette was removed using both I-SceI induction and an additional round of Red-mediated recombination according to the following procedure. One mL LB cultures of *E. coli* GS1783 containing candidate Kan^R^ recombinants were grown at 32°C to logarithmic phase (OD_600_ = 0.6), followed by dilution with 1 mL of prewarmed LB plus 2% L-arabinose for an additional hour to induce I-SceI cleavage. These cultures were then shifted to a 42°C shaking water bath for 30 min to activate Red-mediated recombination, shifted back to 32°C for 2-3 hrs of recovery (∼OD_600_ = 0.5), plated on LB with 34 μg/mL chloramphenicol and 1% L-arabinose, and incubated at 32°C overnight. Single colonies were then screened for sensitivity to kanamycin and tested using the PCR strategy described above. Finally, scarless introduction of the single *nsp5* C**<u>C</u>**C to C**<u>A</u>**C mutation (M^pro^ P132H) was confirmed in RA-SARS2-M^pro^-H132 BAC by BAC DNA preparation (Qiagen Large-Construct Kit, 12462) and full-BAC sequencing was performed using large plasmid sequencing (Plasmidsaurus).

### Cell culture and virus production

Vero-E6-TMPRSS2-T2A-ACE2 (Vero-E6-AT) cells were obtained from BEI Resources (NR-54970) and 293T cells from ATCC (CRL-3216). The Caco2-ACE2 (Caco2-A) cells were made in-house and originated from Caco2 (ATCC, HTB-37). The A549-ACE2-DPP4-TMPRSS2 (A549-ADT) cells were kindly provided by the Perlin lab at the Center for Discovery and Innovation, Hackensack Meridian Health, Nutley, NJ, USA (30). Unless indicated, cell lines were maintained in Dulbecco-modified Eagle’s medium (DMEM, high glucose, Thermo Fisher Scientific, 11965092) supplemented with 10% fetal bovine serum (FBS, Biowest, 058N24), 100 U/mL penicillin, and 100 μg/mL streptomycin (Thermo Fisher Scientific, 15140122). Caco2-A cells were maintained in Eagle’s minimum essential medium (EMEM, ATCC, 30-2003) supplemented with 10% FBS. Cells were cultured at 37^◦^C in a 5% CO_2_ atmosphere with humidity. Cells were routinely tested for mycoplasma contamination using a MycoAlert® Mycoplasma Detection Kit (Lonza, LT07-318).

Primary human bronchial epithelial cells (HAE) were obtained from Epithelix (EP51AB) and expanded in PneumaCult™-NGEx Medium (STEMCELL Technologies, 100-1505). Upon reaching confluence, cells were detached using Gibco™ TrypLE™ Express Enzyme (1×) (Thermo Fisher Scientific, 12604013) and seeded onto permeable ThinCert™ cell culture inserts (Greiner Bio-One, 662641). PneumaCult™-NGEx medium was added to both the apical and basolateral compartments until the cells reached confluence. The apical medium was then removed to establish an air-liquid interface (ALI), and the basolateral medium was replaced with PneumaCult™-ALI-S Medium (STEMCELL Technologies, 05050). Mucus was removed every other day by washing the apical surface with PBS. HAE cultures were differentiated for 4 weeks to generate fully differentiated, polarized pseudostratified mucociliary epithelium resembling the native human airway epithelium. Cultures were maintained at 37°C in a humidified incubator with 5% CO₂. The reference SARS-CoV-2 strain USA-WA1/2020 (WT SARS2) was reported previously (9, 17). Virus stocks were generated by infecting confluent monolayers of Vero-E6-AT cells and incubating for 48 h at 37°C in 5% CO₂. Virus-containing medium was harvested, clarified by centrifugation, aliquoted, and stored at −80°C. Mock stocks were prepared in parallel from uninfected cells maintained under identical conditions. Stock titers were determined by plaque assay.

RA-SARS2 viral stocks were generated by transfecting semi-confluent Vero-E6-AT cells in 6-well plates with 4 µg of RA-SARS2-M^pro^-H132 BAC DNA using lipofectamine 2000 (Thermo Fisher Scientific, 11668019). At 24 hours post-transfection, the medium was replaced with DMEM containing 10% FBS, 100 U/mL penicillin, and 100 μg/mL streptomycin. After an additional 24 hours (48 hours post-infection, hpi), the infected cells were transferred to T-75 flasks and fresh growth medium was added. When virus-induced cytopathic effect (CPE) was observed, as evident by syncytia formation (∼48 hpi) and cell lysis (∼72-96 hpi), viral supernatants were collected, filtered using a Millex-HV PVDF 0.45 µm filter (Sigma, SLHVR33RB), and stored at -80°C (P0 stock of RA-SARS2). Viral stocks were passaged/expanded by adding 1 mL of prior viral stock with 14 mL of DMEM supplemented with 2% FBS and 100 units/mL penicillin/streptomycin (represented as inoculation medium in the next sections) onto confluent Vero-E6-AT cells in T-75 flasks. As stated above, when severe CPE was evident, the viral supernatants were harvest and either stored or used for experimentation. Viral RNA was then extracted from the viral supernatants using the Quick-RNA Viral Kit (Zymo Research, R1035) and sent for RNA-seq for verification (Plasmidsaurus).

### Virus sequencing

Raw reads were trimmed with fastp (v0.23.2) and aligned to the SARS2 reference genome (GenBank accession MN985325.1) using Bowtie2 (v2.4.4). Depth and coverage were calculated with SAMtools (v1.16.1) and visualized in R (v4.4.3) to confirm the deletion of ORF3a or ORF7a and substitution in the M^pro^ sequence at the P132 amino acid position.

### Immunofluorescent microscopy

Vero-E6-AT cells were plated into chambered microscope slides (Falcon culture slides, Corning, 354114), grown for 24 hours, and infected with viral supernatant (2000 PFU/ well). After an additional 48-hour incubation at 37°C, the infected cells were fixed with 4% paraformaldehyde (Thermo Fisher Scientific, 047392.9M) and permeabilized with PBS 0.5% Triton X-100 (Sigma, T9284-500mL). Non-specific antigens were blocked with PBS containing 0.5% Triton X-100 and 2.5% goat serum (Fisher Scientific, 16-210-07) and then slides were incubated for 1 hour with anti-SARS2 spike antibody (Genetex, GTX632604) diluted 1:300 in blocking solution. Slides were then washed 3 times with blocking solution and incubated for an additional hour with secondary antibody (Goat anti-Mouse IgG (H+L) Highly Cross-Adsorbed Secondary Antibody, Alexa Fluor Plus 488, Invitrogen, A32723) diluted 1:500 in blocking solution. The slides were washed 3 more times with PBS, stained with Hoechst 33342 using the manufacturer’s protocol (Thermo Fisher Scientific, H3570), and mounted with ProLong Glass Antifade Mountant (Invitrogen, P36980). The samples were visualized using a Zeiss confocal microscope (LSM710), and the images were analyzed using Fiji software (ImageJ2 Version 2.16.0/1.54p).

### Viral plaque assays

Confluent monolayers of Vero-E6-AT cells were inoculated with 10-fold serial dilutions of viral stocks in inoculation medium. After ∼1.5 hpi, cells were covered with 1.5% carboxymethyl cellulose (Millipore Sigma, C4888-500G) and incubated for 5 days at 37°C. After incubation, the cells were fixed using a final concentration of 10% neutral buffered formalin (NBF, Azer Scientific, 20NBF-4-G), stained with 0.5% crystal violet solution (Thermo Fisher Scientific, C581-25), and washed with water. Plates were then air-dried and imaged using a LI-COR Biosciences Odyssey imaging system and the plaque forming units (PFU)/mL for each reaction was calculated.

### Cytopathic effect (CPE) assays

Confluent monolayers of Vero-E6-AT cells in 96-well plates were treated with various antiviral compounds using 5-fold serial dilutions across 10 points, starting with an initial concentration of 20 µM. ESV (HY-143216), NMV (HY-138687), and TQR (HY-10550) were purchased from MedChem Express, while RTV (SML0491) was obtained from Sigma-Aldrich. Virus controls (VC) and cell controls (CC) were maintained and tested in parallel. Compounds were added onto cells individually or in combination 3 hours prior to infection with RA-SARS2 (120 PFU/well). After 4 dpi, the supernatant was removed, CellTiter-Glo 2.0 reagent (Promega, G9242) was added to the cells, shaken for 10 minutes, and then luminescence was measured using a Tecan Spark plate reader. Percent viral inhibition was calculated using the following formula:

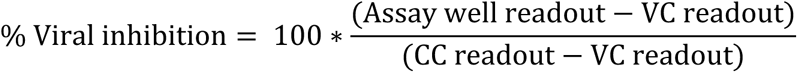

Percent viral infection was then determined by the following equation:

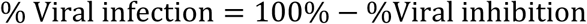

Additionally, CPE was also determined by crystal violet staining. To inactivate the virus and fix the cells for crystal violet staining, 20% NBF was added to a final concentration of 10%. The plates were then washed with water and then stained with 0.5% crystal violet. Finally, the plates were again washed with water, air-dried, and imaged using a LI-COR Biosciences Odyssey imaging system.

### Reduced sensitivity mutant production and sequencing

Confluent monolayers of Vero-E6-AT cells in 12-well plates were treated for 3 hours with 25 nM ESV (∼1/2 of ESV EC_50_ in this system) and then infected with 3200 PFU of RA-SARS2. Multiple independent VCs were maintained in parallel. From 1 to 5 dpi, the cells were observed for CPE and the cell supernatants (filtered as above) were individually stored in -80°C, avoiding any cross-contamination. For the next passage, confluent monolayers were similarly infected with stored viral culture supernatant at a 1:5 ratio with increasing concentrations of ESV with each passage (ESV concentration increased 2-fold with each forward passage, however the concentration was kept the same when viruses displayed sensitivity). After the collection of all viral supernatants, the plates were fixed, stained with crystal violet, and imaged to determine overall CPE. Finally, reduced-sensitivity mutants were grown in the presence of 500 nM of compound for maintenance. RNA was extracted as above, cDNA was produced using the Transcriptor High Fidelity cDNA Synthesis Kit according to the manufacturer’s protocol (Roche, 5081963001), and Phusion™ High-Fidelity DNA Polymerase was used for amplification of the *nsp5* gene using the following primers: 5’-GTGGAGCAATGGATACAACTAGC (forward) and 5’-TCATTGCAAAAGCAGACATAGCA (reverse). PCR amplicons were then cleaned-up/gel-purified (Monarch Spin PCR and DNA Cleanup Kit, New England Biolabs, T1130L), quantified by nanodrop, and subjected to Sanger sequencing using the amplification primers above (Eurofins Genomics).

### Viral Infection for RNA quantification

Vero-E6-AT, Caco-2A, and A549-ADT cells were seeded in 96-well plates and, upon reaching 100% confluency, infected with RA and WT SARS2 using an inoculum of 40 PFU/well (Vero-E6-AT) or using an inoculum of 400 PFU/well (Caco-2A and A549-ADT). Three wells were dedicated to each time point. At 2 h post-infection, the inoculum was removed, cells were washed twice with PBS, and fresh medium was added. At 24, 48, 72, and 96 hpi, 100 µL of supernatant was collected from each of the three wells assigned to that timepoint into an equal volume of DNA/RNA Shield (Zymo Research, R110050), and the corresponding monolayers were lysed in 100 µL DNA/RNA Shield for 5 min and harvested. Collected samples were stored at −80°C prior to viral RNA isolation.

HAE cultures were apically infected with RA- and WT-SARS2 (50,000 PFU/well) or mock infected for 2 h at 37°C, as described (31). Virus and mock inocula were diluted in PBS, and 50 μL was added to the apical surface of each insert. Following infection, cultures were washed three times with PBS, and the third PBS wash was collected as the 2 hpi sample. The cultures were subsequently maintained at the air-liquid interface for the remainder of the experiment.

Apical samples were collected every 24 h until 96 hpi. For each collection, 100 μL of PBS was added to the apical surface of each insert and incubated for 10 min at 37°C. The apical wash was then collected into DNA/RNA Shield and stored at −20°C for subsequent viral RNA quantification by RT-qPCR.

### Isolation of nucleic acids and quantitative PCR

Viral RNA was extracted from cell culture supernatants and cells using the Quick-RNA Viral Kit (Zymo Research, R1035) according to the manufacturer’s protocol. Viral RNA levels were determined by RT-qPCR targeting the nucleocapsid (*N*) gene, using the GoTaq Probe 1-Step RT-qPCR System (Promega, A6120) on a LightCycler 480 II instrument (Roche) with primers and a probe as described (32) [forward primer 5ʹ-CACATTGGCACCCGCAATC (600 nM), reverse primer 5ʹ-GAGGAACGAGAAGAGGCTTG (800 nM), and probe 5ʹ-FAM-ACTTCCTCAAGGAACAACATTGCCA-BHQ1 (200 nM)] (32). Reverse transcription was performed at 45 °C for 15 min, followed by 2 min at 95 °C and 40 cycles of 15 s at 95 °C and 1 min at 56 °C. Absolute N gene copy numbers were determined against a plasmid standard curve. The N gene amplicon was cloned into pJET1.2 using the CloneJET PCR Cloning Kit (Thermo Fisher Scientific, K1231), and the resulting plasmid was linearized with HindIII, purified with the GeneJET PCR Purification Kit (Thermo Fisher Scientific, K0701), and quantified by Nanodrop. Copy number was calculated from the length of the linearized plasmid (3104 bp) and an average nucleotide mass of 320 g/mol using Avogadro’s constant, and seven 10-fold serial dilutions of the linearized plasmid were used as standards.

### Gain-of-signal assay for M^pro^ activity in living cells

For a luciferase-based gain-of-signal assay for M^pro^ activity in living cells, 3×10^6^ 293T cells were seeded in a 10-cm dish and transfected after 24 hours with a mixture containing 2 µg of the Src-SARS2-M^pro^-Tat-fLuc construct (33) and 6 µl of TRANS-IT-LT1 transfection reagent (Mirus, MIR2304). Mutant M^pro^ constructs with single amino acid substitutions were generated via site-directed mutagenesis using Phusion HF DNA Polymerase (NEB, M0530L) following the manufacturer’s protocol and the following primer sets (D48G: 5’-GTGAGGGTATGCTTAATCCCAAT and 5’-AGCATACCCTCACTAGTGCAGAT; L89F: 5’-CTGAAGTTCAAAGTCGATACTGCA and 5’-CTTTGAACTTCAGTACGCAATTCT; M49L: 5’-GAGGATCTGCTTAATCCCAATTAC and 5’-TAAGCAGATCCTCACTAGTGCAG). All constructs underwent verification by Sanger sequencing. Four hours post-transfection, cells were washed once with PBS, trypsinized, resuspended, and counted. Cells were then diluted to a concentration of 4×10^5^ cells/mL, and 50 µL of this suspension was plated into each well of a 96-well plate containing 50 µL of media with the desired drug concentration, resulting in a 1× final drug concentration and 2×10^4^ cells per well. At 48 hours post-transfection, 50 µL of Bright-Glo reagent (Promega, E2610) was added to each well, followed by a 2-minute incubation, after which luminescence was measured using a Tecan Spark plate reader.

M^pro^ activity at each compound concentration was calculated relative to DMSO-treated transfected controls, which were defined as 100% activity. Raw luminescence values (RLU) from compound-treated cells were normalized to the mean RLU of DMSO-treated transfected cells, and percent activity was calculated as follows:

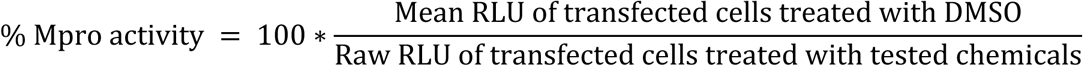

### Biochemical M^pro^ activity assays

An established biochemical assay was used to quantify M^pro^ and mutant derivative cleavage activity *in vitro* (33–37). Fluorescence is liberated by M^pro^ cleavage between Q and S of a peptide substrate, DABCYL-KTSAVLQ|SGFRKM-EDANS (UPBio, V1010-1). Cleavage reactions were carried out in 20 µL reactions in 384-well plates (Greiner Bio-One, 781074) with 5 µM substrate, 50 nM M^pro^, 20 mM Tris-HCl, pH 8.0, 150 mM NaCl, 1mM EDTA, 0.05% Tween20, 0.1 mg/mL bovine serum albumin (BSA), 1 mM DTT. For inhibition studies, M^pro^ was incubated at room temperature with various concentrations of chemical (2-fold serial dilution series starting at 100 µM) for 30 min in reaction buffer prior to addition of the substrate to initiate the reaction. Fluorescence intensity was monitored kinetically using a Tecan Spark multimode plate reader (Tecan Life Sciences) (Ex. 355 nm / Em. 538 nm). Initial reaction velocities were determined by fitting a linear regression to the initial linear phase (t=0-30 min) of each fluorescence progress curve and calculating the slope (ΔRFU/Δt) using Prism.

To determine the percent activity of M^pro^ for a compound concentration series, test well reaction velocity was normalized to the mean initial velocity of vehicle (DMSO treated wells):

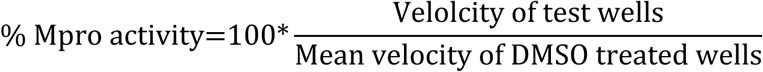

### NanoDSF thermal-shift assays

The thermal stability of recombinant SARS2 M^pro^ WT and mutant proteins (D48G, M49L, L89F, and M49L/L89F) was assessed using nano differential scanning fluorimetry (nanoDSF). Purified proteins were diluted to a final concentration of 0.1 mg/mL in buffer containing 20 mM Tris-HCl (pH 8.0), 100 mM NaCl, and 0.5 mM TCEP. Protein samples (10 µL each) were loaded into Prometheus High Sensitivity Capillaries (NanoTemper Technologies, PR-C006) according to the manufacturer’s instructions, with care taken to avoid the introduction of air bubbles or particulate matter. Thermal unfolding was monitored using a Prometheus Panta instrument (NanoTemper Technologies). Intrinsic tryptophan and tyrosine fluorescence was excited at 280 nm, and fluorescence emission was recorded at 330 and 350 nm. Samples were heated from 35 °C to 85 °C at a ramp rate of 5 °C/min. Fluorescence intensities at 330 and 350 nm were recorded throughout the thermal ramp. The fluorescence ratio (F330/F350) was plotted as a function of temperature using GraphPad Prism. Each protein condition was analyzed in three technical replicates.

## RESULTS

### Introduction of a precise mutation into SARS2 M^pro^ using *En Passant* mutagenesis

One of the most safe and versatile reverse genetics systems for SARS2 studies is a BAC-based system harboring the WA1 strain full-length genome, with attenuating deletions in *orf3a* and *orf7b* (18, 38) (**Fig. 1A**). Deletion of accessory ORFs does not abolish SARS2 replication in permissive cell lines (*e.g*., Vero-E6-AT), but substantially reduces replication efficiency and viral fitness in more immune-competent models (*e.g*., Caco-2A, A549-ADT cells and primary human airway epithelial cultures, HAE), and attenuates infection *in vivo* in murine and hamster models (18, 21, 39). However, a drawback to this model system is that the WA1 and other early coronavirus lineages are effectively extinct from present day human populations, and any variants accumulated since the original COVID-19 pandemic such as those in current Omicron lineage viruses are not represented.

**Fig. 1.**
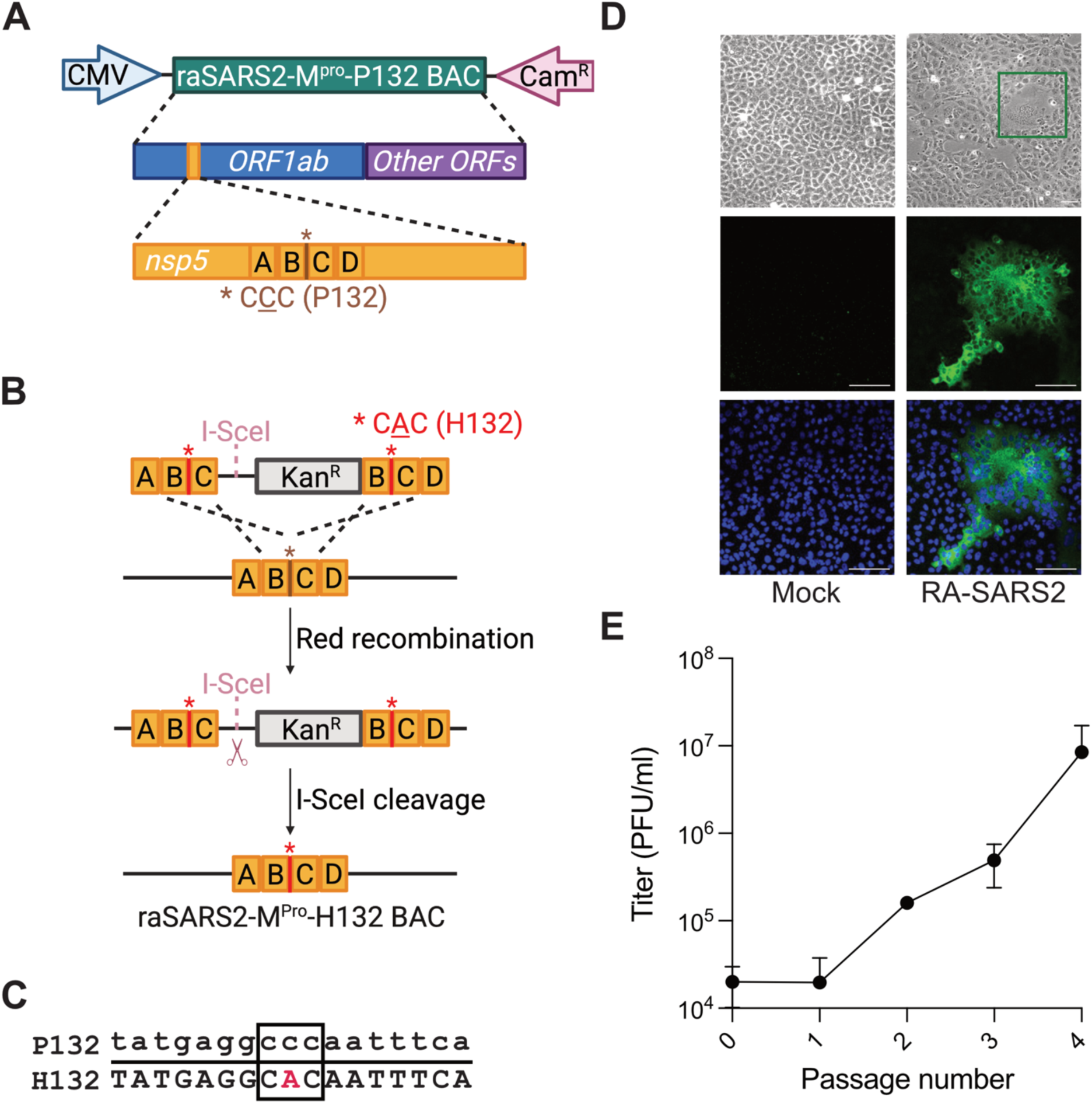
Production of RA-SARS2 with Omicron M^pro^. **(A)** Schematic of the RA-SARS2-M^pro^-P132 BAC. Regions A-B-C-D represent homology used for *En Passant* mutagenesis. **(B)** Schematic of the steps required for *En Passant* mutagenesis and scarless replacement of *nsp5* C**<u>C</u>**C codon (P132) with a C**<u>A</u>**C codon (H132) to generate RA-SARS2-M^pro^-H132 BAC. See text for details. **(C)** Sequence analysis of RT-PCR amplicon confirmed the *nsp5* C**<u>C</u>**C (P132) to C**<u>A</u>**C (H132) codon substitution (nucleotide sequence highlighted by a black box). (**D)** Brightfield images of Vero-E6-AT cells at 48 hpi demonstrating CPE, including syncytia formation, in RA-SARS2 infected cultures but not in mock-treated controls (scale bar = 100 µm). Immunostaining at 48 hpi showed viral S protein expression (green) in RA-SARS2 infected Vero-E6-AT cells (scale bar = 100 µm; nuclei stained blue with Hoechst). **(E)** Serial passage of RA-SARS2 supernatants on fresh Vero-E6-AT cells results in increased viral titers (mean PFU/mL +/- SD of n = 3 technical replicates).

To mitigate this issue for the viral main protease, M^pro^, a recombineering approach called *En Passant* mutagenesis (28) was used to replace the single WA1 remanent amino acid P132 (C**<u>C</u>**C codon) with the current Omicron variant amino acid H132 (C**<u>A</u>**C) (**Fig. 1B**). Briefly, high-fidelity PCR was used to generate a kanamycin-resistance (Kan^R^) cassette flanked by 60 nucleotides of sequence homologous to the M^pro^ open-reading-frame *nsp5*. The homologous sequence is identical apart from the single base substitution mutation (C-to-A) that is required to change codon 132. The cassette was then transformed into *E. coli* containing the RA-SARS2-M^pro^-P132 BAC, and Kan^R^ clones were identified by locus-specific PCR and DNA sequencing. Next, Kan^R^ clones were streak-purified on L-arabinose containing plates to induce I-SceI expression and double-stranded DNA cleavage adjacent to the Kan^R^ cassette, which is subsequently repaired by looping-out recombination between homologous regions (B-C) flanking the Kan^R^ cassette. The resulting RA-SARS2-M^pro^-H132 BAC was screened for kanamycin sensitivity (Kan^S^), and locus-specific PCR and DNA sequencing was used to verify seamless deletion of the drug resistance cassette and incorporation of the C**<u>A</u>**C codon for H132 (**Fig. 1C**).

Next, RA-SARS2-M^pro^-H132 BAC DNA was transfected into Vero-E6-AT cells to recover a viral stock, RA-SARS2. Virus functionality was demonstrated by a visible CPE, including syncytia formation and cell death, at 48 hours post-transfection. After observing prominent CPE and cell death, the viral supernatants were collected and used for infection on fresh Vero-E6-AT cells. The infected cells were then assessed by immunostaining for viral S protein (**Fig. 1D**). Cell-free viral supernatants were serially passaged on fresh Vero-E6-AT cultures to propagate the virus and to generate a high-titer RA-SARS2 stock as quantified by viral plaque assays (**Fig. 1E**).

### Antiviral drug testing with RA-SARS2

Prior work using the original RA-SARS2 system demonstrated that this virus is sensitive to individual drugs including ESV and Remdesivir (19). To confirm and extend this work, EC_50_ values were determined for RA-SARS2 treated with ESV and NMV. These values were determined using Vero-E6-AT cells in a quantitative CPE assay utilizing CellTiter-Glo 2.0, and a qualitative cell viability assay using crystal violet staining. Consistent with prior literature (4), we found that the ESV EC_50_ is 45 nM (**Fig. 2A**) and that the NMV EC_50_ is 1128 nM (**Fig. 2B**) in the quantitative CPE assay. These results are reflected visually in the qualitative cell viability assay data using crystal violet staining (**Fig. 2C**).

**Fig. 2.**
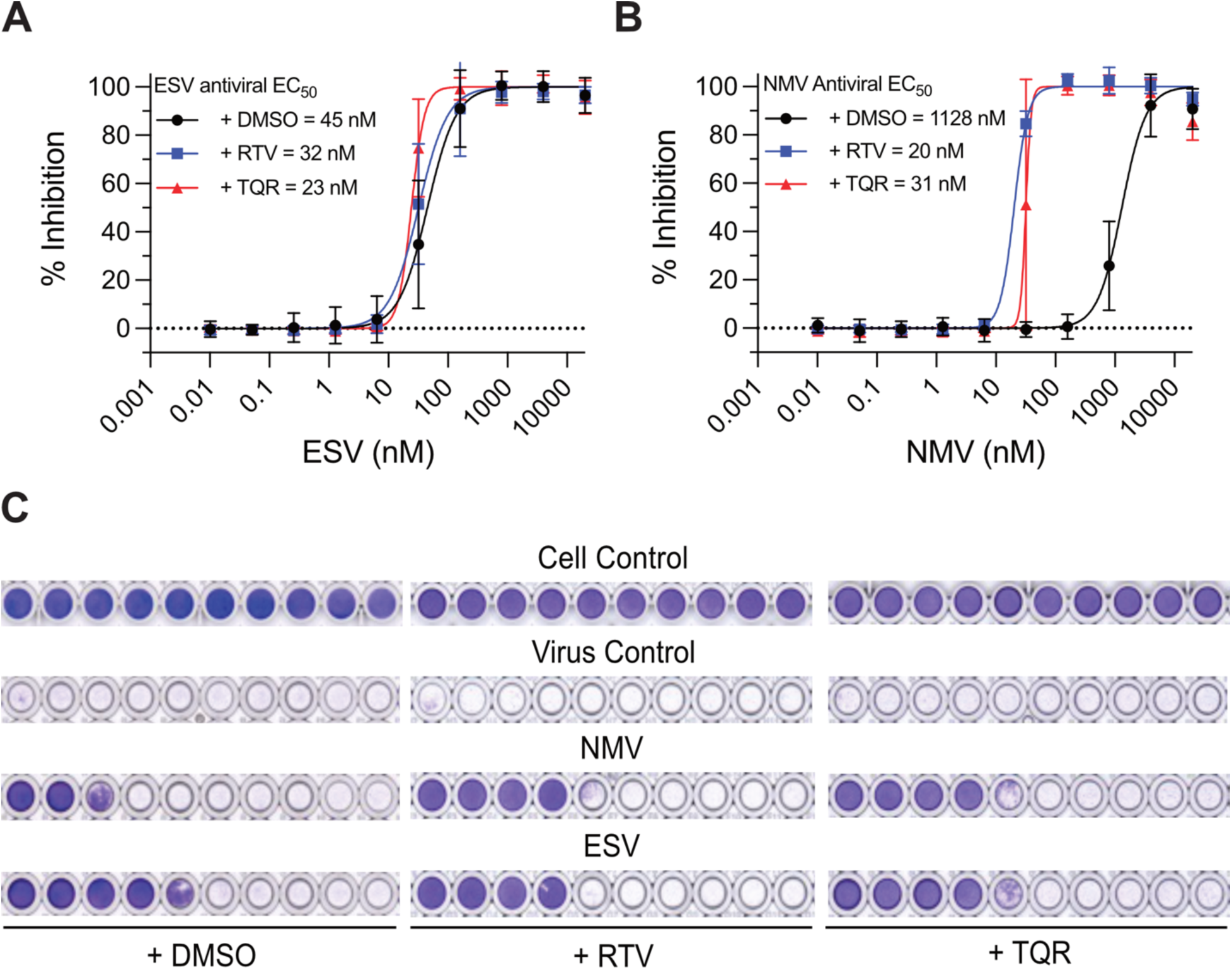
Activity of antiviral compounds ENS and NMV. (**A-B**) Dose response curves showing EC_50_ values for ESV and NMV, respectively, in the presence of RTV (blue), TQR (red), or control DMSO (black). Individual data points are from at least n = 3 biologically independent experiments +/- SD of CPE read-outs. (**C**) Representative images of crystal violet-stained Vero-E6-AT cells following treatment with the indicated dose ranges of NMV and ESV in the presence of DMSO, RTV, or TQR.

The low potency of NMV in the Vero-E6-AT system is likely due to cellular cytochrome P450 activity (Cyp3A4) and/or to compound efflux by the P-Glycoprotein (P-Gp) transporter (3). To distinguish between these possibilities, dose response experiments were conducted in the presence of 10 µM RTV, which inhibits both Cyp3A4 and P-Gp, or 1 µM TQR, which is a selective P-Gp blocking agent (40–43). The addition of RTV and TQR had little effect on the antiviral EC_50_ of ESV, as expected (**Fig. 2A**, **2C**). In contrast, both compounds dramatically enhanced the potency of NMV, with EC_50_ values now considerably lower at 20 nM and 31 nM, respectively (**Fig. 2B**, **2C**). These results demonstrate the utility of this system for not only measuring antiviral activity of a compound but also determining whether the compound is susceptible to cellular efflux through P-Gp.

### Assessment of RA-SARS2 permissiveness in human cell lines

RA-SARS2 was subsequently tested in a panel of human infection models alongside Vero-E6-AT cells, an African green monkey kidney cell line that remains the most permissive model for SARS2 but does not reflect the human tissue context of infection. The human models included A549-ADT cells as a receptor-enhanced respiratory epithelial system in which susceptibility depends on the addition of viral entry factors, Caco-2A cells as a permissive intestinal epithelial system with ACE2 receptor expression, and primary human airway epithelial (HAE) cultures as a physiologically relevant *ex vivo* airway system. These models allowed for comparison of viral infection across systems that vary in species and tissue origin, entry factor expression, and innate antiviral responses. Wildtype SARS-CoV-2 WA1 (WT SARS2) was used as the non-attenuated parental control, since RA-SARS2 was derived from the WA1 strain. Both WT and RA-SARS2 virus stocks were titrated on Vero-E6-AT by plaque assay prior to infection experiments.

Visible CPE was first evaluated in the cell lines by crystal violet staining (**Fig. 3A**). The clearest CPE for RA-SARS2 was observed in Vero-E6-AT cells, followed by Caco-2A cells. No observable CPE was detected in A549-ADT cells infected with RA-SARS2. In Vero-E6-AT cells, both RA and WT SARS2 established robust infection. Although CPE was reduced for RA-SARS2 relative to WT SARS2, it remained detectable down to 26 PFU for RA-SARS2, compared with 0.4 PFU for WT SARS2. Consequently, crystal violet staining was most informative for comparing CPE in Vero-E6-AT cells, whereas CPE assessment in human cell lines was less straightforward. Although Caco-2A cells supported infection and showed a comparable reduction in CPE for RA-SARS2 relative to WT SARS2, the virus-induced cytopathic changes were overall less pronounced than in Vero-E6-AT cells, limiting the utility of this model for direct CPE comparison. In A549-ADT cells, CPE was evident following WT SARS2 infection but entirely absent across all RA-SARS2 doses tested, indicating that the lack of cytopathic changes was specific to the attenuated virus.

**Fig. 3.**
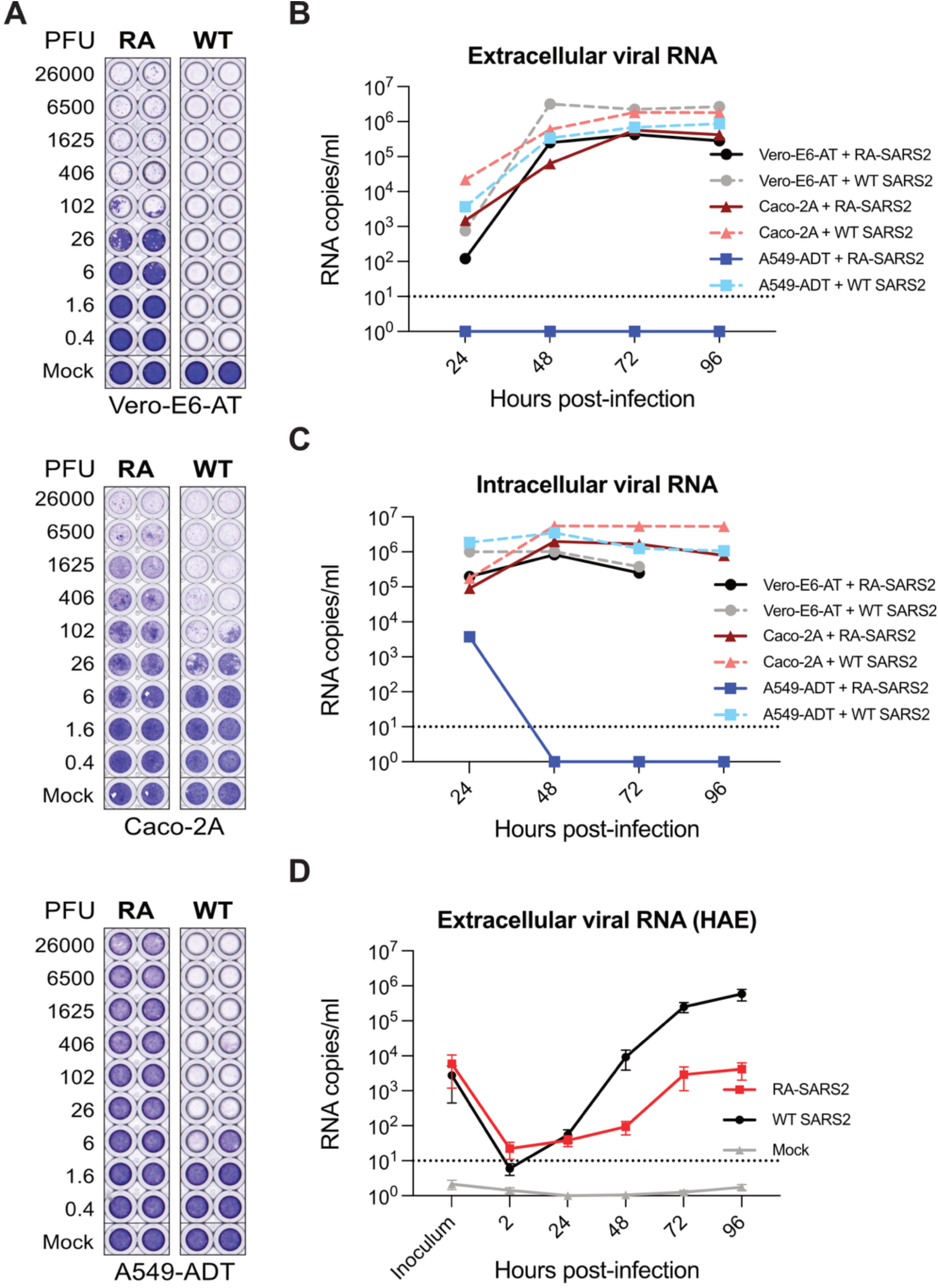
Cellular model systems indicate safety of RA-SARS2 system for antiviral research. (**A**) Crystal violet staining of Vero-E6-AT, Caco-2A, and A549-ADT cells infected with serial 4-fold dilutions of RA-SARS2 (RA) or WT SARS2 (WT), ranging from 26,000 to 0.4 PFU/well, alongside mock-infected controls. Loss of staining indicates virus-induced CPE. Wells are shown in duplicate. All experiments were performed in at least two biological replicates. (**B-C**) Replication kinetics of RA-SARS2 and WT SARS2 determined by RT-qPCR of extracellular and intracellular viral RNA, respectively. Vero-E6-AT cells were infected with an inoculum value of 40 PFU/well and Caco-2A and A549-ADT cells with 400 PFU/well. Supernatants and cell lysates were collected every 24 hpi and viral RNA quantified as copies/mL. Values below the limit of detection are plotted at 1 copy/mL. Intracellular RNA is not shown for Vero-E6-AT cells at 96 hpi, as complete lysis of the monolayer precluded reliable lysate collection. Each experiment was performed in two biological replicates, each with at least three technical replicates (each data point is the mean ± SD). (**D**) Replication of RA-SARS2 (red) and WT SARS2 (black) in HAE cultures, with mock-infected cultures included as a control (grey). Apical washes were collected at the indicated time points and extracellular viral RNA quantified by RT-qPCR as copies/mL. Each experiment was performed in at least three biological replicates, each with at least two technical replicates (each data point is the mean ± SD).

As permissivity does not always match the severity of visible CPE, infection was also monitored using RT-qPCR. To compare the replication kinetics of RA and WT SARS2 within each model, Vero-E6-AT cells were infected with an inoculum of 40 PFU/well, whereas Caco-2A and A549-ADT cells were infected with an inoculum of 400 PFU/well, and supernatants and cell lysates were collected every 24 hours post-infection. Viral RNA in culture supernatants was measured to assess extracellular viral RNA levels (**Fig. 3B**), whereas analysis of cell lysates was used to evaluate intracellular viral RNA accumulation (**Fig. 3C**). Extracellular RNA levels closely mirrored the crystal violet staining results: in Vero-E6-AT and Caco-2A cells, RA-SARS2 RNA levels were approximately 1 log_10_ lower than those of WT SARS2, whereas no RA virus replication was detected in A549-ADT cells. Interestingly, intracellular RNA levels differed minimally between RA and WT SARS2 in Vero-E6-AT and Caco-2 cells. Taken together with the extracellular data, these results suggest a modest reduction in the release of viral particles during RA-SARS2 infection in these two cell lines. Additionally, RA-SARS2 RNA was detected in A549-ADT cells only at 24 hpi and was undetectable at all subsequent time points, indicating that although this cell line is susceptible to RA-SARS2 entry, it does not support productive replication. In Vero-E6-AT cells, both RA and WT SARS2 infected cells efficiently, leading to complete cell lysis by 96 hours post-infection even at the lower viral inoculum amount. Consequently, intracellular RNA could not be reliably measured by RT-qPCR at this time point in Vero-E6-AT cells, but extracellular RNA, which was already released into the supernatant, remained detectable. Virus replication was also examined in the *ex vivo* HAE model by RT-qPCR (**Fig. 3D**). WT SARS2 displayed robust replication kinetics in HAE cultures, reaching maximal viral RNA levels at 96 hpi, whereas RA-SARS2 replication was reduced by approximately 100-fold.

### RA-SARS2 M^pro^-M49L exhibits reduced sensitivity to ESV

To model the acquisition of reduced antiviral sensitivity in a safe live-virus system, RA-SARS2 was serially passaged in Vero-E6-AT cells under increasing ESV selection pressure, beginning at 25 nM and increasing 2-fold in concentration at each passage, up to a final concentration of 4 µM. At each passage, virus-containing supernatants were transferred onto fresh cells maintained with the compound; passages were terminated at 5 dpi, and supernatants were carried forward to the next passage even when CPE was limited.

Reduced ESV sensitivity became apparent at approximately after the 5^th^ passage in a subset of cultures. RT-PCR and Sanger sequencing of the M^pro^ coding region in the ESV-selected populations revealed the characteristic M49L substitution in three ESV-selected isolates and a D48G substitution in one; two of the M49L isolates additionally acquired an L89F substitution. In comparison, no changes were found in control (CT) RA-SARS2 passaged in parallel in DMSO. A representative chromatogram illustrating an ATG>CTT (M49L) change relative to the DMSO-passaged control is shown in **Fig. 4A**. Of note, one M49L lineage transiently displayed an M49I intermediate at passage 5 (ATG>ATT>CTT), suggesting that M49L may be the more stable of the two ESV-selected substitutions at this position.

**Fig. 4.**
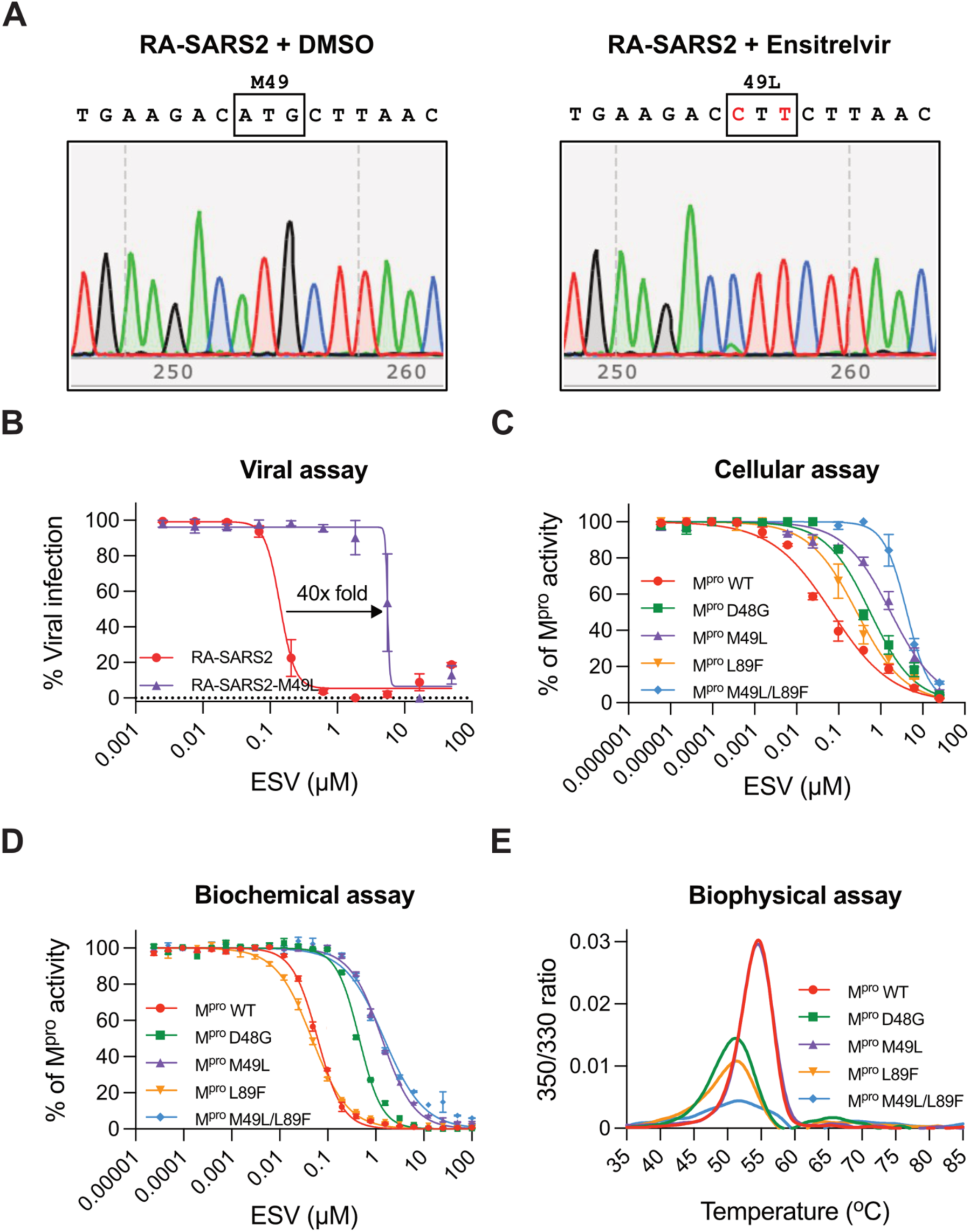
The RA-SARS2 system provides insights into ESV-reduced sensitivity mutants. **(A)** M^pro^ coding region sequence of RA-SARS2 populations passaged in the presence of DMSO (control, left) or ensitrelvir (ESV, right). **(B)** Antiviral susceptibility of RA-SARS2 encoding the M49L substitution (purple) compared with the DMSO-passaged control virus (CT RA-SARS2, red), determined in dose-response assays across the indicated range of ESV concentrations. Viral infection is expressed as a percentage of the untreated control. Each experiment was done at least twice (biological replicates) with at least two technical replicates (mean ± SD). **(C)** Cellular gain-of-signal (GoS) assay measuring ESV inhibition of WT M^pro^ (red), D48G (green), M49L (purple), L89F (orange), and M49L/L89F (blue). M^pro^ activity is expressed as a percentage of the untreated control for condition. Each experiment was done at least twice (biological replicates) with at least two technical replicates (mean ± SD). **(D)** Biochemical assay measuring ESV inhibition of WT M^pro^ and the indicated mutants (color code as in panel C). M^pro^ activity is expressed as a percentage of the uninhibited enzyme. Each experiment was done at least twice (biological replicates) with at least two technical replicates (mean ± SD). **(E)** Thermal unfolding profiles by nanoDSF of WT M^pro^ and the indicated mutants (color code as in panel C). Profiles are representative of two biologically independent experiments.

As M49L is the most frequently reported substitution in circulating ESV-resistant strains and was the most prevalent change in our selection procedure, it was chosen as a representative mutant for evaluation in the live-virus context. Antiviral susceptibility of the CT RA-SARS2 and the M49L RA-SARS2 variant was assessed in dose-response assays with ESV (**Fig. 4B**). The M49L variant displayed a pronounced rightward shift in its ESV dose-response curve relative to CT RA-SARS2, amounting to an approximately 40-fold increase in EC_50_. Both viruses were fully inhibited at the highest concentrations tested, indicating a shift in potency rather than complete loss of ESV activity.

M^pro^ WT, M^pro^-M49L, and the additional amino acid substitution variants were characterized in cellular (**Fig. 4C**), biochemical (**Fig. 4D**), and biophysical (**Fig. 4E**) assays. These orthogonal systems are particularly informative for resistance studies, as they allow multiple M^pro^ amino substitutions to be compared in parallel and help to distinguish direct effects on inhibitor binding from indirect effects on M^pro^ folding and stability.

To determine whether the substitutions directly impact M^pro^ inhibitor sensitivity, all four variants (D48G, M49L, L89F, and M49L-L89F) were evaluated alongside WT M^pro^ in a previously described (33) cellular gain-of-signal (GoS) assay for M^pro^ activity (**Fig. 4C**). All substitutions conferred reduced ESV sensitivity relative to WT M^pro^, with the rank order L89F < D48G < M49L < M49L/L89F. The M49L-L89F double substitution produced the largest rightward shift in the dose-response curve, supporting a direct role for both substitutions in altering ENS sensitivity.

The effects of the substitutions were further examined using biochemical assays with purified WT M^pro^ (**Fig. 4D**). In this system, L89F alone displayed inhibition comparable to WT M^pro^, whereas D48G conferred an intermediate shift and M49L and M49L-L89F conferred the largest reductions in ESV potency. The difference between the cellular and biochemical assays for L89F suggests that this substitution contributes to reduced sensitivity primarily in a cellular context rather than by directly altering inhibitor binding to the isolated enzyme.

Finally, these M^pro^ mutant proteins were analyzed using nano-differential scanning fluorimetry (nanoDSF), a biophysical method to quantify the thermal stability of purified proteins (**Fig. 4E**). WT and M49L M^pro^ displayed nearly identical thermal transitions, indicating that M49L reduces ESV susceptibility without destabilizing the protein. In comparison, D48G, L89F, and M49L-L89F showed transitions at lower temperatures and with markedly reduced amplitude, consistent with decreased thermal stability of these variants. These measurements provided additional insights into each amino acid substitution’s impact on M^pro^ at the protein level and enabled comparisons in the absence of a cellular environment. Collectively, the live-virus, cellular, biochemical, and biophysical data indicate that the identified substitutions reduce susceptibility to ESV in the RA-SARS2 model, with M49L as the dominant contributor and L89F acting as a secondary, context-dependent modifier.

## DISCUSSION

*En passant* mutagenesis is a straightforward and efficient method for modifying RNA virus genomes, enabling seamless edits without leaving residual sequences that might impact viral fitness. When used with SARS2, this technique provides a practical way to generate changes in attenuated viruses, which are safer to handle than fully replication-competent strains. Attenuated viruses lessen the biosafety concerns linked to fully replication-competent viruses, which may facilitate antiviral testing in various experimental settings. Attenuated viruses are especially useful for studying amino acid substitutions that confer a reduced sensitivity to drugs such as ESV as well as candidate novel antiviral compounds.

An important consideration when working with attenuated viruses is that attenuation alters replication capacity in a cell-type-dependent manner. In our hands, RA-SARS2 replicated efficiently in Vero-E6-AT and Caco-2A cells, with only a modest reduction relative to WT SARS2, but did not establish productive replication in A549-ADT cells and was markedly restricted in HAE cultures. The degree of attenuation therefore cannot be assumed to be uniform across models, and permissiveness must be established empirically in each system before use. This constrains the applicability of attenuated viruses to questions that depend on faithful reproduction of the virus-host relationship, including studies of innate immune antagonism, tropism, and pathogenesis, where the altered replication phenotype confounds interpretation of the cellular response.

For antiviral susceptibility testing, however, these constraints are far less restrictive. Provided a permissive cell system is selected, standard readouts transfer directly to the attenuated virus: CPE scoring and crystal violet staining remain fully applicable and can be complemented by RT-qPCR quantification of viral RNA where cytopathic changes are less pronounced. Moreover, serial passage of RA-SARS2 under ESV selection recovered the M49L substitution that predominates among ESV-selected and circulating resistant strains, together with D48G and L89F, and the resulting shift in susceptibility was consistent with reported values for the corresponding substitutions in wildtype virus. The attenuated system therefore yields drug sensitivity data that may foretell clinical outcomes, while operating within a considerably safer experimental framework.

Finally, information generated in the viral context can be substantially strengthened by orthogonal M^pro^ assays. A cellular gain-of-signal assay, as well as biochemical and biophysical assays, allow multiple substitutions to be compared in parallel and, importantly, distinguish between effects on inhibitor potency, catalytic activity, and protein stability, a resolution that live-virus assays alone cannot provide. Together, attenuated live-virus studies and orthogonal target-based assays constitute a tractable and safety-conscious workflow for characterizing antiviral resistance mechanisms.

## Acknowledgments

We thank Michael Carpenter, Tony Lin, and present Harris lab members for helpful feedback. This work was supported by National Institute of Allergy and Infectious Disease (NIAID) grants U19-AI171954 and U19-AI171403. Partial salary support for RD was provided by NIAID 1F31AI189116-01. RSH is an Investigator of the Howard Hughes Medical Institute and the Ewing Halsell President’s Council Distinguished Chair at the University of Texas at San Antonio.

